# Metabolism-focused CRISPR screen unveils Mitochondrial Pyruvate Carrier 1 as a critical driver for PARP inhibitor resistance in lung cancer

**DOI:** 10.1101/2023.11.29.569226

**Authors:** Takashi Furusawa, Renzo Cavero, Yue Liu, Haojian Li, Xia Xu, Thorkell Andresson, William Reinhold, Olivia White, Myriem Boufraqech, Thomas J. Meyer, Oliver Hartmann, Markus E. Diefenbacher, Yves Pommier, Urbain Weyemi

## Abstract

Homologous recombination (HR) and poly ADP-ribosylation are partially redundant pathways for repair of DNA damage in normal and cancer cells. In cell lines that are deficient in HR, inhibition of poly (ADP-ribose) polymerase (PARP1/2) is a proven target with several PARP inhibitors (PARPi) currently in clinical use. Resistance to PARP inhibitors often develops, usually involving genetic alterations in DNA repair signaling cascades, but also metabolic rewiring particularly in HR-proficient cells. PARP1/2 utilize NAD+ (nicotinamide adenine dinucleotide), an essential substrate not only for PARPs but also for multiple pathways in cellular metabolism, TCA cycle and mitochondrial functions. Thus, NAD+ is central to many key cellular functions. Both activation of PARPs by DNA damage and their inhibition by drugs such as olaparib affect NAD+ consumption. We surmised that alterations in NAD+ metabolism by cancer drugs such as Olaparib might be involved in the development of resistance to drug therapy. To test this hypothesis, we conducted a metabolism-focused CRISPR knockout (KO) screen to identify genes which undergo alterations during treatment of tumor cells with PARP inhibitors. Of about 3000 genes in the screen, our data revealed that mitochondrial pyruvate carrier 1 (MPC1) is an essential factor in desensitizing NSCLC lung cancer lines to PARP inhibition. In contrast to NSCLC lung cancer cells, triple negative breast cancer cells do not exhibit such desensitization following MPC1 loss and reprogram the TCA cycle and oxidative phosphorylation pathways to overcome PARP inhibitor treatment. Our findings unveil a previously unknown synergistic response between MPC1 loss and PARP inhibition in lung cancer cells.

## Introduction

Modulating DNA repair efficiency via genetic or biochemical means can be harnessed to sensitize cancer cells to chemotherapy and to identify new players in genomic stability. For example, homologous recombination (HR) and poly ADP-ribosylation (PARylation) are two partially redundant DNA repair pathways, the latter anchored by poly (ADP-ribose) polymerases (PARP1/2) which are triggered by DNA damage to utilize NAD^+^ (nicotinamide adenine dinucleotide) to poly ADP-ribosylate themselves and other target proteins ^1–4^. However, NAD^+^ is central to energy metabolism, a coenzyme for redox reactions, and an essential cofactor for non-redox NAD^+^-dependent enzymes, including sirtuins, CD38 as well as PARP1/2. Thus, NAD^+^ can influence many key cellular functions. NAD^+^ consumption may lead to activation of other pathways in attempts to replenish the NAD^+^ pool ^3^.

Several PARP1/2 inhibitors (PARPi) have been found to be clinically effective in HR-deficient cancers ^5, 6^, but the treatment often triggers prosurvival responses particularly in HR-proficient cancer cells ^7^. In this study, we surmise that, given the significant role of PARP family members as metabolic sensors and NAD^+^ consumers, PARPi resistance may reflect an intrinsic association between PARP-dependent DNA repair and NAD^+^-mediated mitochondrial energetic reprogramming. This primary hypothesis is based on mounting evidence linking DNA damage signaling to metabolic pathways, including mitochondrial respiration, glycolysis, the pentose phosphate pathway, and redox homeostasis ^2, 8–11^. To test this hypothesis, we utilized a metabolism-centered CRISPR-Cas9 genetic screen ^12^ in MDA-MB-231 cells treated with the PARP inhibitor, Olaparib. Our data unveiled mitochondrial pyruvate carrier 1 (MPC1) as a key modulator of resistance to PARP inhibition. We found that MPC1 loss robustly sensitizes lung and breast cancer cells to PARP inhibition *in vitro*. However, breast cancer cells exhibit strong resistance to PARP inhibitors *in vivo*, presumably via metabolic rewiring. Indeed, our data revealed that, unlike a transient silencing, a permanent deletion of MPC1 in triple negative breast cancer cells led to a robust activation of mitochondrial oxidative phosphorylation and activation of TCA cycle upon PARP inhibition. Taken together, our study reveals a novel therapeutic option for targeting PARP in lung cancer cells, while identifying a putative pathway whereby breast cancer cells resist PARP inhibition.

## MATERIALS AND METHODS

### Cell Culture and Reagents

The human breast cancer cell line MDA-MB-231 (from American Type Culture Collection (ATCC)) was grown at 37 °C with 5% CO_2_ in Dulbecco’s Modified Eagle Medium (DMEM) (HyClone), supplemented with 10% Fetal Bovine Serum (Gemini Bio). The mouse breast tumor model cell line 4T1-Luc2 (from American Type Culture Collection (ATCC)) was grown at 37 °C with 5% CO_2_ in Dulbecco’s Modified Eagle Medium (DMEM) (HyClone), supplemented with 10% Fetal Bovine Serum (Gemini Bio). The human non-small cell lung cancer (NSCLC) cell line NCI-H1299 (from Division of Cancer Treatment and Diagnosis (DCTD) Tumor Repository, NCI) was grown at 37 °C with 5% CO_2_ in Roswell Park Memorial Institute (RPMI) 1640 Medium (HyClone), supplemented with 10% Fetal Bovine Serum (Thermo Fisher Scientific) and 1% sodium pyruvate 100mM (Thermo Fisher Scientific). The mouse NSCLC model cell line KP5 (provided by Dr. Markus E. Diefenbacher)^13^ was grown at 37 °C with 5% CO_2_ in Dulbecco’s Modified Eagle Medium (DMEM) (HyClone), supplemented with 5% Fetal Bovine Serum (Gemini Bio). Cells were authenticated using a colorimetric signal amplification system and tested for mycoplasma contamination (R&D systems). All media were supplemented with penicillin and streptomycin (Gibco).

Silencer Select siRNAs were purchased from Thermo Fisher Scientific. Assay IDs for each gene are as follows: *MPC1* (s28488), *ACSM4* (s226320 and s50838), *SLC5A7* (s34076), *PLA2G7* (s1549), and *CLCN7* (s3149). siRNA was transfected by Lipofectamin RNAiMAX (Invitrogen) according to the manufacturer’s instructions. Cell viability was determined by CellTiter-Glo Luminescent Cell Viability Assay (Promega) or PlestoBlue Cell Viability Reagent (Thermo Fisher Scientific) according to the manufacturer’s instructions. PARP inhibitor Olaparib was purchased from Selleckchem (S1060). Olaparib was dissolved in DMSO for stock solution.

### Mouse studies

*NOD scid gamma (NSG)* mice were provided from NCI-Frederick. All animal experiments complied with the protocols for animal use, treatment, and euthanasia approved by the National Cancer Institute Institutional Animal Care and Use Committees.

### Metabolism-Centered CRISPR/Cas9 KO Library Screen

In this study, the human CRISPR metabolic gene library was used to identify metabolic genes responsible for PARP-inhibition resistance in the breast cancer cell line, MDA-MB-231. The library was a gift from David Sabatini’s laboratory to Addgene (*Addgene* #110066). Briefly, we transduced the library which contains 29,698 gRNAs targeting 2,981 human metabolic genes (∼10 gRNAs per gene and 499 control gRNAs targeting intergenic region) at a low MOI (∼0.3) to ensure effective barcoding of individual cells. Then, the transduced cells were selected with 1 μg mL^−1^ of puromycin for 7 days to generate a mutant cell pool, which were then split into three groups. One group was frozen and designated as Day 0 sample. The other two groups were treated with vehicle (DMSO) and Olaparib (2.5 μM) for 14 days, respectively. After treatment, at least 16 million cells were collected for genomic DNA extraction to ensure over 500X coverage of the human CRISPR metabolic gene library. The sgRNA sequences were amplified using NEBNext^®^High-Fidelity 2X PCR Master Mix and subjected to Next Generation Sequencing by the Genomic Sequencing and Analysis Facility of the University of Texas at Austin. The sgRNA read count and hits calling were analyzed using the *MAGeCKFlute* pipeline. Read counts for the CRISPR Screen are shown in <u>Dataset S1</u>.

### RNA-seq and transcriptomics

Single guide RNA (sgRNA)-mediated knockdown was generated in MDA-MB-231 cells for mitochondrial pyruvate carrier 1 (sgMPC1). MDA-MB-231 cells were split into four conditions including sgCTRL and sgMPC1 treated with vehicle (DMSO), along with sgCTRL, and sgMPC1 treated with Olaparib (10 μM). Cells were incubated for 24 days in DMEM before treatment with Olaparib for 5 days. The drug was replenished every two days for the duration of the experiment.

Cells were lysed and processed for RNA using the RNeasy Mini Plus RNA extraction kit (Qiagen). Samples were processed using NuGEN’s Ovation RNA-Seq System V2 and Ultralow V2 Library System and sequenced on an Illumina HiSeq 2500 machine as 2×125nt paired-end reads.

Raw FASTQ files were processed using the RENEE RNA-sequencing pipeline (https://github.com/NCIPangea/RENEE). In brief, Cutadapt (v1.18) was used to trim reads for adapters and low-quality bases. Star v2.5 was then used in 2-pass mode to align the trimmed reads to the human reference genome (hg38). Next, expression was quantified using RSEM v1.3.0. Downstream analysis and visualization were performed within the NIH Integrated Data Analysis Platform (NIDAP) using R programs developed on the Foundry platform (Palantir Technologies). Genes were filtered for low counts (<1 cpm), and quantile normalized prior to differential expression using limma voom v3.38.3. Gene set enrichment analysis (GSEA) was performed using fGSEA v.1.8.0. Differentially expressed genes (p<0.01, FC>2) were further analyzed and pathways with an adjusted p<0.001 were considered significant in the enrichment. Online database Human Mitocarta 3.0 was provided by the Broad Institute.

### Antibodies

Immunoblots were performed using rabbit anti-MPC1 antibody (1:1000, Cell Signaling Technology 14462S) and rabbit anti-GAPDH antibody (1:2500, Cell Signaling Technology 3683S). Secondary antibody HRP-linked rabbit-IgG was used from Cell Signaling Technologies (cat# 9559).

### Viral transduction

To generate MPC1 knockout cells, MDA-MB-231 cells and NCI-H1299 cells were infected with pLentiCRISPR V2 viral vector (Addgene, # 52961) in which the sgRNA for human *MPC1* gene (5’-AAGTCTCCAGAGATTATCAG-3’) was cloned. To generate MPC1-knockdown cells, KP5 cells and 4T1 cells were infected with lentiviral particles produced using shRNA expressing plasmids (pLKO.1) targeting MPC1. The shRNA sequence used is listed as follows: shMPC1: 5’-CAAACGAAGTAGCTCAGCTCA-3’.

### Mouse experiments

All animals were treated in accordance with the recommendations of the NIH Animal Care and Use Committee (ACUC). All animal procedures were performed according to protocols approved by NCI Laboratory Animal Sciences Program (LASP). Intravenous (IV) injection (MDA-MB-231 cells and 4T1 cells) and subcutaneous (SC) injection (KP5 cells) were performed as previously described ^8^. For IV injection, six- to eight-week-old female immunocompromised NOD SCID gamma (NSG) mice (provided from NCI-Frederick) were injected with cells via the lateral tail vein using 29-gauge needles and followed up for metastases burden. In brief, 1 × 10^6^ cells suspended in 200 µL DMEM were injected into the tail vein of each mouse on Day 0. After tumors became established in the lung on Day 1, mice were randomized and treated with Olaparib (50 mg/kg) by oral gavage. To visualize lung metastasized tumors, mice were injected D-luciferin (Gold BioTechnology) at a dose of 150 mg/kg in PBS IP injection and anesthetized with 3–5% isofluorane by inhalation prior to imaging. Images were acquired by Xenogen IVIS Lumina system (Caliper Life Sciences).

For SC injection, six- to eight-week-old female NSG mice were injected with cells at the right flank using 29-gauge needles and followed up for tumor burden. In brief, 350,000 cells suspended in 200 µL DMEM supplemented 50% Matrigel (Corning) were injected into the right flank of each mouse on Day 0. After tumors became established on the skin and tumor volume become 50-100mm^3^, mice were randomized and treated with Olaparib at the dose of 50 mg/kg. Tumor was measured its length (L) and width (W), and the tumor volume (mm^3^) was estimated by the following formula: 1/2 x L(mm) x W (mm) x W (mm). Mice were euthanized when the tumor volume exceeded 2,000 mm^3^ or length exceeded 20mm. Mice (six to eight per treatment group) received the following agents by oral gavage as specified by the experimental protocols: vehicle (10% w/v DMSO, 10% w/v 2-Hydroxypropyl-β-cyclodextrin) or Olaparib (50 mg/kg) in vehicle, daily for 7 days a week. After 3 weeks, animals were sacrificed and examined macroscopically and microscopically for the presence of metastases.

### Reversed-Phase Ion-Pairing LC-MS^2^ Assay for Measuring Cell Central Carbon Metabolites

All reference target compounds (CCM) were purchased from Sigma-Aldrich (St. Louise, MO) (Table 1). The stable isotope labeled internal standards (SI-CCM) were ^13^C_3_-lactate, ^13^C_4_-succinic acid, obtained from Cambridge Isotope Laboratory (Andover, MA) as well as ^13^C_6_-glucose-6-phosphate and ^13^C_6_-fructose-1,6-diphosphate purchased from Medical Isotopes, Inc. (Pelham, NH). All CCM and SI-CCM analytical standards have reported chemical and isotopic purity ≥ 98%. They were used without further purification. OmniSolv^®^ LC-MS grade acetonitrile and methanol were obtained from EMD Millipore (Billerica, MA). Tributylamine (TBA), LC-MS grade acetic acid and formic acid were purchased from Fisher Scientific (Hampton, NH). All chemicals and solvents used in this study were HPLC or reagent grade unless otherwise noted.

For cell CCM assay, 500 µL chilled 80% methanol-water solution was added to the cell pellet as previously described ^14^. Sample was vortexed vigorously for 30 sec and centrifuged at 14,000 g for 10 min. Fifty µL supernatant was transferred to an autosampler vial containing 50 µL 10 µM SI-CCM methanol solution. Sample was dried with the SpeedVac^®^ vacuum concentrator (ThermoFisher Scientific, Waltham, MA) and then reconstituted in 60 µL 3% (v/v) methanol in water. Ten µL sample was injected for reversed-phase ion-pairing LC-MS^2^ analysis. Reversed-phase ion-pairing LC-MS^2^ analysis was performed using a Thermo TSQ™ Quantiva triple quadrupole mass spectrometer (Thermo Scientific, San Jose, CA) coupled with a NexeraXR LC system (Shimadzu Scientific Instruments, Columbia, MD). Both the HPLC and mass spectrometer were controlled by Xcalibur™ software (Thermo Scientific). Reversed-phase ion-pairing liquid chromatography was carried out on a 100-mm long x 2.1-mm i.d. Synergi Hydro-RP C18 column with 2.5 µm particles and 100 Å pore size (Phenomenex, Torrance, CA) and kept in 40 °C. The mobile phase, operating at a flow rate of 200 µL/min, consisted of 10 mM TBAA in water as solvent A and methanol as solvent B. For the analysis of CCM and SI-CCM, a linear gradient stayed at B/A solvent ratio 3:97 for 3 min, then changed the B/A solvent ratio from 3:97 to 80:20 in 14 min. After washing with 98% B for 3 min, the column was re-equilibrated with a mobile phase composition B/A of 3:97 for 10 min prior to the next injection. The general MS conditions were as follows: source: ESI; ion polarity: negative; spray voltage: 2500 V; sheath and auxiliary gas: nitrogen; sheath gas pressure: 40 arbitrary units; auxiliary gas pressure: 5 arbitrary units; ion transfer capillary temperature, 350 °C; scan type: selected reaction monitoring (SRM); collision gas: argon; collision gas pressure: 2 mTorr. Quantitation of cell CCM was carried out using Xcalibur™ Quan Browser (Thermo Scientific). Calibration curves for each CCM were constructed by plotting CCM/SI-CCM peak area ratios obtained from calibration standards versus CCM concentrations and fitting these data using linear regression with 1/*X* weighting. The CCM concentrations in samples were then interpolated using this linear function.

### Oxygen consumption rate (OCR) measurement using Seahorse Analyzer

Metabolic measurements were carried out in standard 96-well Seahorse microplates on a Seahorse XF24 analyzer. Pyruvate oxidation was measured using oxygen consumption rate (OCR) when cells were incubated in unbuffered Seahorse media containing 10 mM sodium pyruvate as the only respiratory substrate. For all experiments 20,000 cells per well were plated 16–18 hours prior to analysis.

### Statistical Methods

Statistical analyses were performed using *Graphpad Prism 9*. Unless otherwise noted, data were analyzed by Student’s t-test and considered significant at p < 0.05.

## Results and discussion

### Metabolism-focused CRISPR screen reveals *mitochondrial pyruvate carrier 1* as a key driver for resistance to PARP inhibitor

To determine whether PARylation in cancer cells fuels metabolic and mitochondrial bioenergetic reprogramming (Figure 1A), we utilized a metabolism-centered CRISPR-Cas9 genetic screen ^12^ in MDA-MB-231 breast cancer cells treated with the PARP inhibitor, olaparib, to identify metabolic genes whose loss enhances cell death upon PARP inhibition (Figure 1B). We utilized the *MAGeCK-MLE* pipeline to assess the degree to which these metabolic genes are required for survival upon PARP inhibition ^15^. The data mining based on the most ranked genes required for resistance to PARP inhibitor revealed Phosphatase and Tensin Homolog (PTEN) as potentially the most required gene for resistance upon treatment with PARP inhibitor Olaparib (Figure 1C). However, when comparing the beta scores for each of the sgRNA guides for PTEN, we found that most guides score positively in both Dimethyl sulfoxide (DMSO) and Olaparib-treated samples, thus pointing to PTEN as a gene likely regulating cell proliferation rather than resistance to PARP inhibitor (Supplementary Figure 1A). To screen for essential metabolic genes required for resistance to PARP inhibitor, we then assess distribution of the sgRNA guides of the top 20 genes putatively required for resistance to PARP inhibition in our screen (Figure 1D). We found that most sgRNA guides for MPC1, ACSM4, PLA2G7, SLC5A7, and CLCN7 show positive beta scores in cells treated with DMSO, while Olaparib treatment led to a reverse pattern of sgRNA guides distribution for these genes (Supplementary Figure 1B-F). The other putatively required genes exhibit an overall negative sgRNA guides distribution, suggesting these genes may simply be essential for the general survival of cancer cells (Supplementary Figure 1G-T). To further ascertain whether PTEN loss leads to cells being unable to mediate resistance to this PARP inhibitor, we generated cells deficient for PTEN and analyzed their sensitivity to olaparib. As shown in Supplementary Figure 2, PTEN deletion with sgRNA targeting PTEN did not yield a prominent increase in cell death upon olaparib treatment in a clonogenic assay, further ruling out a possibility for PTEN to mediate resistance to PARP inhibition in MDA-MB-231 cells.

**Figure 1.**
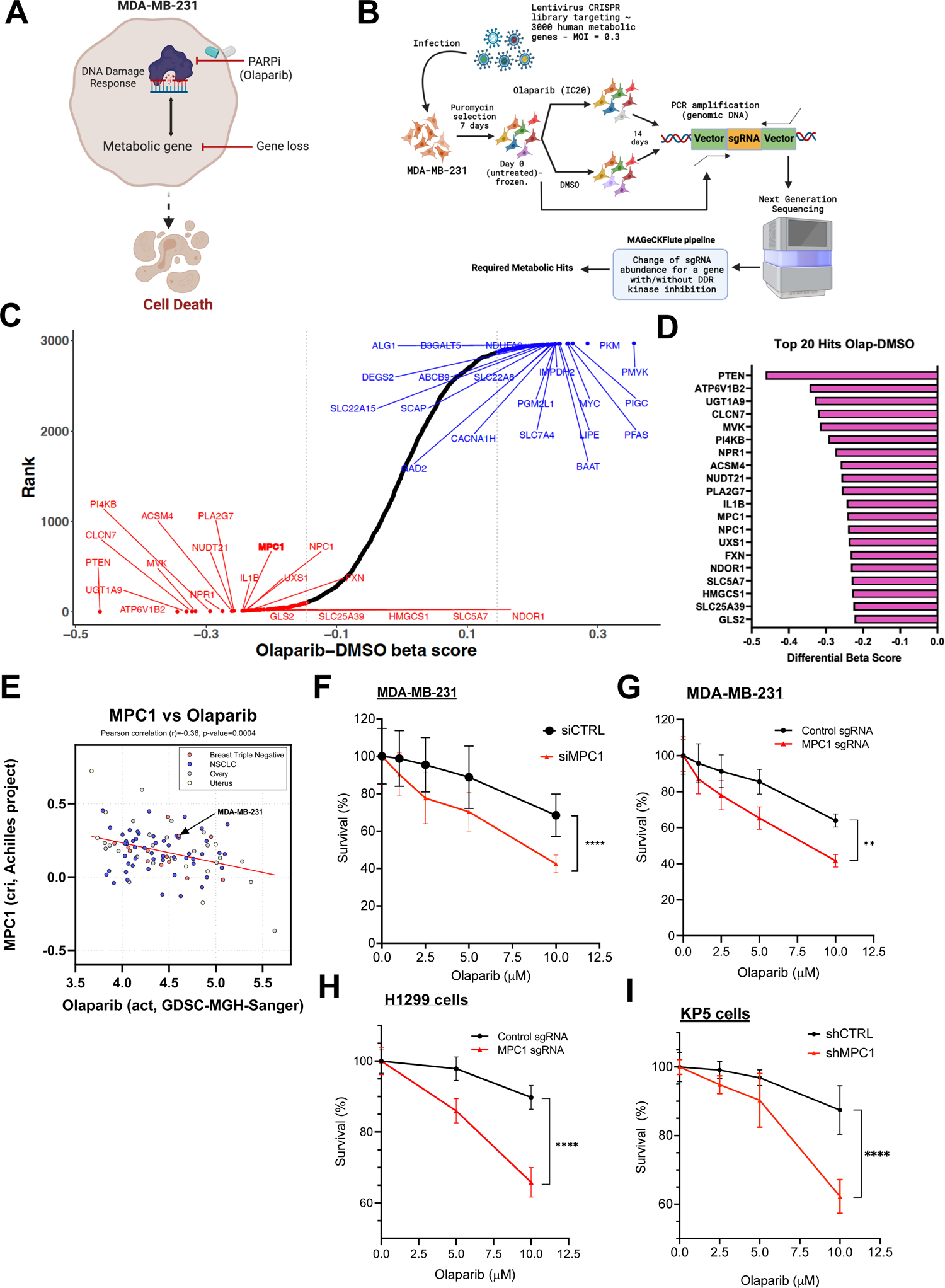
Metabolism-centered CRISPR/Cas9 KO library screen identified MPC1 as a driver for resistance to PARP inhibition. (A) Hypothetical dialogue between the PARP-mediated DNA damage response and other metabolic processes in cancer cells. Olaparib treatment prevents DNA damage response by inhibiting PARP, whose role is to facilitate the localization of DNA repair machinery to damaged DNA sites but in doing so, also consumes its substrate NAD+, altering intracellular NAD+ pool levels. By this means, Olaparib treatment may lead to a metabolic reprogramming through its effects on NAD+ pools. Thus, silencing of a putative metabolic gene, i.e., one not directly involved in DNA repair, responsible for resistance to PARP inhibition may sensitize cancer cells to death. (B) Schematic diagram illustrating the workflow of metabolism centered CRISPR/Cas9-expressing lentiviral vector KO library screen. This screen enables the evaluation of the contribution of ∼2981 metabolic enzymes and metabolism-related transcription factors as well as 500 control sgRNAs to drug resistance as previously described. (C) The Rank plot of genes generated by *MAGeCK-Flute-MLE*, which is sorted based on differential beta score by subtracting the DMSO beta score from the Olaparib beta score. (D) The top 20 genes with the lowest differential beta score. (E) *CRISPR Achilles* dataset were utilized to plot for MPC1 expression levels and the degree of olaparib activity in cancer cells (breast cancer, ovarian cancer, and uterus cancer). Triple negative breast cancer cells are highlighted. (F) Survival of MDA-MB-231 cells after treatment with indicated siRNAs and Olaparib for 6 days. (G) Survival of sgRNAs infected-MDA-MB-231 cells after treatment with indicated Olaparib concentrations for 6 days. Control and MPC1-targeting sgRNAs were used. Note that MDA-MB-231-sgMPC1 cells are considered MPC1 knockout pool cells. These cells have a low residual expression of MPC1. (H) Survival of sgRNA-infected H1299 and (I) shRNA-infected KP5 cells after treatment with indicated Olaparib concentrations for 6 days. Data are represented as mean SD; n = 5. Statistical significance was determined by two-tail unpaired student t test. **P < 0.01; ****P < 0.0001.

To identify the most required gene for resistance to PARP inhibition, we then utilized siRNA screen targeting the five (5) genes bioinformatically ranked as putative resistance genes (CLCN7, ACSM4, PLA2G7, SLC5A7, and MPC1), followed by measurement of cell proliferation upon Olaparib treatment (Supplementary 3A). Our data revealed MPC1 silencing led to a ∼ 50.9% increase in cell death while depletion of CLCN7, ACSM4, PLA2G7, and SLC5A7 led to a similar response to Olaparib treatment as did MDA-MB-231 cells transfected with siRNA control (∼24% to 34%) (Supplementary 3B). Taken together, these observations point to MPC1 as a potential driver of resistance to PARP inhibition in MDA-MB-231 cells.

To investigate whether the relationship between PARP inhibition and MPC1 in tumor cells may provide clues for any biological correlation, we conducted a comparative analysis of MPC1 survival level and olaparib activity in a large panel of breast, ovarian, lung, and uterine cancer cell lines using the *CRISPR Achilles* (*GDSC-MGH-Sanger*) datasets. The data indicate that the transcript levels of MPC1 knockout negatively correlates with enhanced activity for Olaparib (Figure 1E), suggesting that MPC1 loss of function facilitates greater resistance to olaparib treatment.

To elucidate the extent to which MPC1 loss sensitizes cancer cells to PARP inhibition, we generated MDA-MB-231 breast cancer cells depleted for MPC1 using either a transient silencing with small interference RNA (siRNA) method or a permanent deletion with CRISPR-KO-Cas9 approach (Supplementary Figure 4). We observed that MDA-MB-231 cells depleted of MPC1 are highly sensitive to Olaparib (Figure 1F, G). Similar results were obtained using both human and murine non-small cell lung cancer (NSCLC) cells H1299 and KP5 respectively (Figure 1H, I). These data point to MPC1 as a critical factor in desensitizing breast and lung cancer cells to PARP inhibitors *in vitro*.

### Mitochondrial pyruvate carrier 1 depletion sensitizes lung cancer cells to PARP inhibitor *in vivo*

To evaluate the requirement for MPC1 in the resistance to PARP inhibitor, we generated lung cancer cells xenografts in immunocompromised *NSG* mice using both control and murine NSCLC cancer cell line KP5 (K-Ras mutated/p53 deletion) depleted for MPC1 in the presence or the absence of the PARP inhibitor Olaparib (Figure 2A). The KP5 cell line was previously established by Diefenbacher’s group and utilized to model lung adenocarcinoma and response to treatment *in vivo* ^13^. Both control and MPC1-depleted cells were transduced with a Luciferase-expressing vector to monitor tumor growth *in vivo* using bioluminescence. As shown in Figure 2B (top panels), MPC1 depletion led to a 60.3% decrease (*p*=0.009) in tumor growth, as did a treatment with Olaparib in mice inoculated with control cells (Figure 2B, left panels, and C). Most remarkably, MPC1 depletion further sensitizes lung cancer cells to PARP inhibition (Figure 2B, bottom panels, and C), findings which are consistent with a 34.0% decrease (*p*=0.006) in tumor weight of MPC1-depleted cells treated with Olaparib at the endpoint (Figure 2D, E). These data strongly suggest that MPC1 loss significantly fosters a metabolic environment prone to an elevated sensitivity to PARP inhibition, thus impairing the ability of cancer cells to progress *in vivo*. These findings imply that lung cancer cells may utilize MPC1-driven metabolism to resist PARP inhibition.

**Figure 2.**
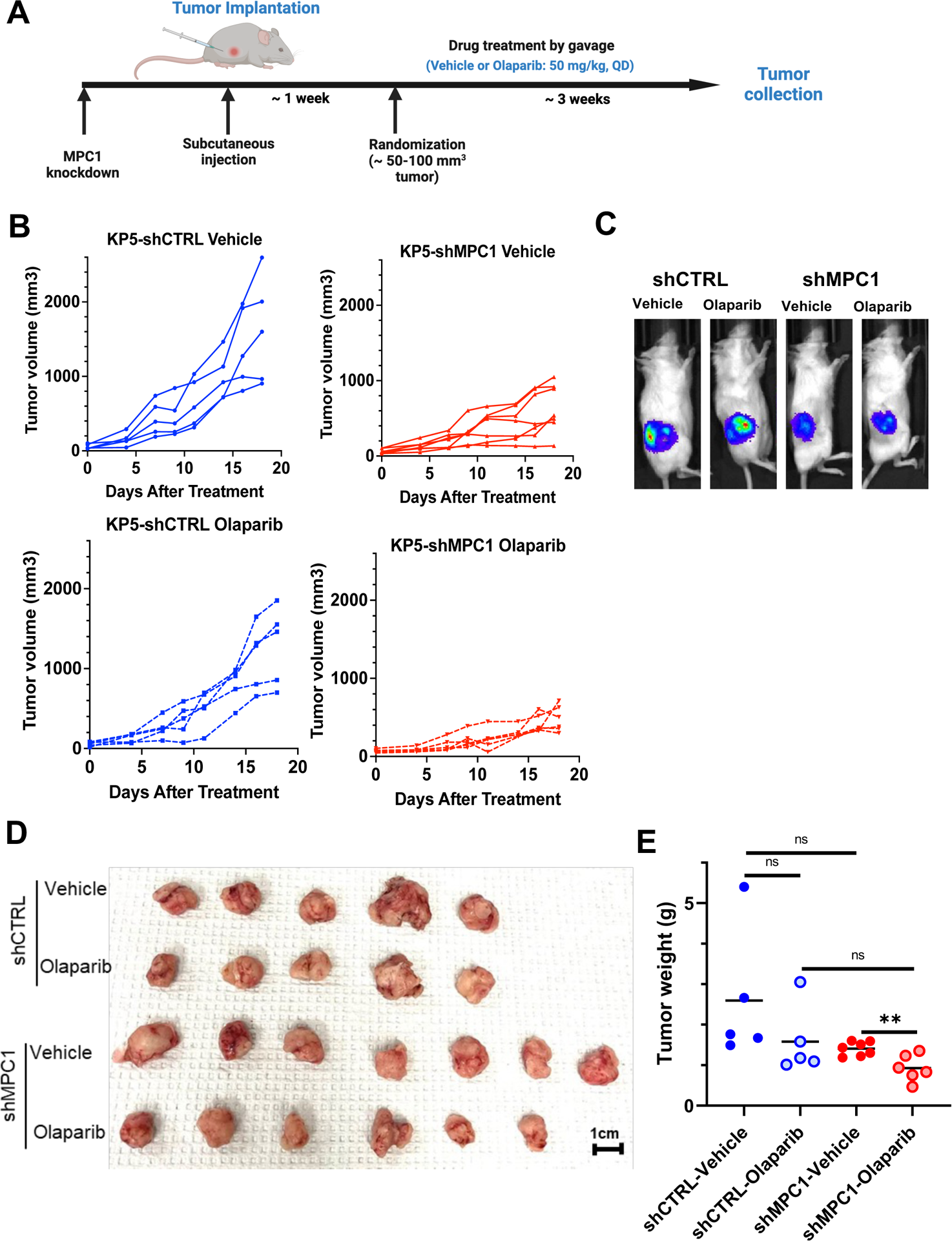
Depletion of MPC1 sensitized mouse NSCLC KP5 cells to Olaparib treatment in immunocompromised NOD SCID gamma mice. (A) Schematic of the experimental tumor model using luciferase expressing-KP5 cells. (B) Measurement of individual tumor volume throughout the experiment. (C) Representative bioluminescence imaging of mice inoculated with KP5-Luc-shCTRL or shMPC1 cells treated with vehicle or PARP inhibitor (Olaparib, 50 mg/kg) at Day18. (D) Image of tumors taken *ex vivo* at Day 18. (E) *Ex vivo* tumor weight at Day 18. Statistical significance was determined by two-tail unpaired student t test. Data are represented as mean SD; n = 5 (shCTRL), n = 7 (shMPC1-Vehicle), n =6 (shMPC1-Olaparib). ns=not significant; **P < 0.01; ***P<0.001.

### Triple negative breast cancer cells robustly reactivate oxidative phosphorylation and TCA cycle to overcome sensitivity to PARP inhibitor *in vivo*

Since our *in vitro* results imply that both lung and breast cells exhibit similar response to PARP inhibition upon MPC1 loss, we set out to determine whether MPC1 deletion may lead to a robust sensitivity of triple negative breast cancer cells to PARP inhibitor *in vivo*. To that end, we generated both human and murine triple negative breast cancer xenografts using MDA-MB-231 and 4T1-Luc2 cells respectively. These control and MPC1-depleted cells were inoculated to immunocompromised *NSG* mice (Figure 3A). Most unexpectedly, we found that MPC1 loss failed to sensitize human triple negative breast cancer MDA-MB-231 xenografts to PARP inhibitor *in vivo* (Figure 3B-D). Similar observations were made in mice inoculated with the murine line 4T1-Luc 2 (Figure 3E, F). We found a modest decrease in tumor growth with MPC1-depleted 4T1-Luc2 cells, findings which were substantiated by 1.72 times extension of median survival rate (*p*<0.0001) in mice inoculated with MPC1-depleted 4T1 cells (Figure 3G). Taken together, our data imply that triple negative breast cancer cells acquire resistance to PARP inhibition, presumably via a metabolic rewiring.

**Figure 3.**
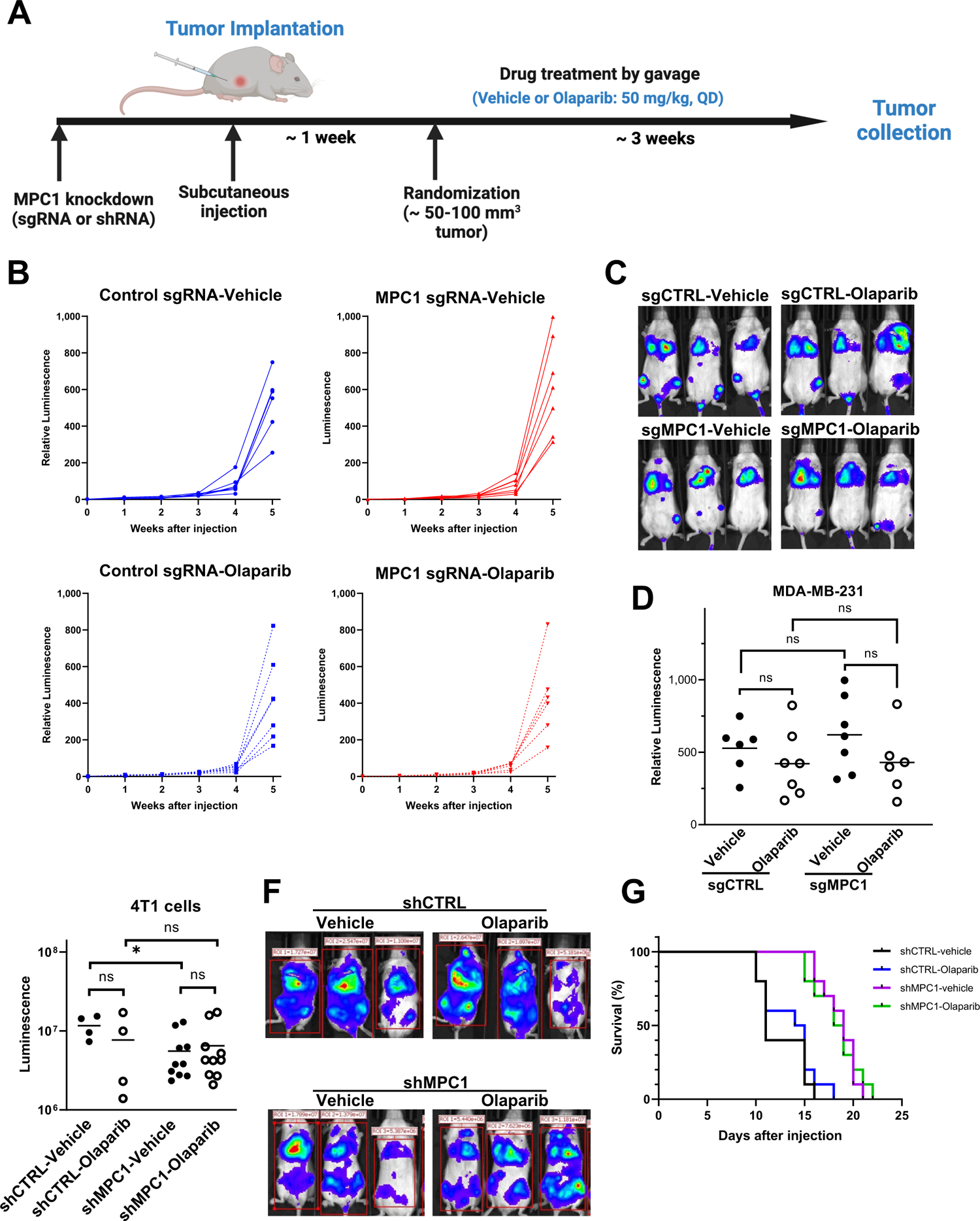
In vivo metastasis assay using MDA-MB231 cells and 4T1 cells. (A) Schematic of the experimental metastasis model using MDA-MB-231-luciferase or 4T1-luciferase cells. (B) Comparison of individual tumor progression in mice with indicated treatment based on the measurement of bioluminescence imaging using IVIS system. The tumor progression is assessed by normalizing the luminescent to Day 0 luminescence. (C) Representative bioluminescence imaging of mice injected with MAD-MB-231-Luciferase sgCTRL or sgMPC1 and treated with vehicle or Olaparib (50 mg/kg) at Week 5. (D) Relative luminescence at week 5. Statistical significance was determined by two-tail unpaired student t-test. Bar represents mean SD; n = 6 or 7. ns=not significant. (E) Measurement of tumors in mice by bioluminescence imaging using IVIS system. Mice are injected with 4T1-luciferase shCTRL or shMPC1 and treated with vehicle or Olaparib (50mg/kg) for 2 Weeks. Statistical significance was determined by two-tail unpaired student t-test. Bar represents mean SD; n = 4 or 10. ns=not significant; *P < 0.05. (F) Representative bioluminescence imaging of mice injected with 4T1-Luciferase shCTRL or shMPC1 treated with vehicle or Olaparib (50 mg/kg) at Week 2. (G) Survival curves for NSG mice injected with 4T1-luciferase cells and treated as indicated. n=10. Statistical significance determined by log-rank test indicates ****P<0.0001 between shCTRL-Vehicle (black) and shMPC1-Vehicle (purple); ***P<0.001 between shCTRL-Olaparib (blue) and shMPC1-Olaparib (green).

To investigate the crosstalk between genomic instability, energetic metabolism, and the potential mechanism of resistance to PARP, we performed an RNA-seq to assess for genome-wide differentially expressed gene (DEG) comparing control and MPC1-depleted MDA-MB-231 cells, following PARP inhibition. We then utilized a Gene Set Enrichment Analysis (GSEA) to further assess the expression pattern of genes involved in energetic metabolism comparing control and MPC1-depleted MDA-MB-231 cells treated with Olaparib for 5 days. Our data revealed a robustly selective activation of oxidative phosphorylation (OXPHOS) pathway in MPC1-depleted cells treated with Olaparib (Figure 4A, B). We then set out to evaluate the degree to which a transient silencing of MPC1 with small interference RNA (siRNA) or its permanent depletion with CRISPR/Cas9 affects mitochondrial respiratory activity using a *Seahorse* assay. As shown in Figure 4C, a transient depletion of MPC1 led to a robust decrease in oxygen consumption rate as revealed by a 30 to 50% reduction in ATP production and maximal respiration respectively (Figure 4D and E), findings which reflect a prominent role of MPC1 in mitochondrial metabolism and respiration ^16, 17^. Olaparib treatment yielded a similar reduction, but not a synergistic response upon MPC1 depletion. A permanent deletion of MPC1 resulted in a reversal of the maximal respiration pattern, with PARP inhibition promoting a 15-20% increase in both ATP production and maximal respiration upon MPC1 loss (Figure 4F-H). Together, these findings imply a mitochondrial energetic rewiring reflecting the reactivation of oxidative phosphorylation following PARP inhibition in breast cancer cells upon permanent deletion of MPC1. Inversely, similar experiments in lung cancer cells KP5 demonstrated a lack of metabolic rewiring upon PARP inhibition in MPC1-depleted cells (Supplementary Figure 5).

**Figure 4.**
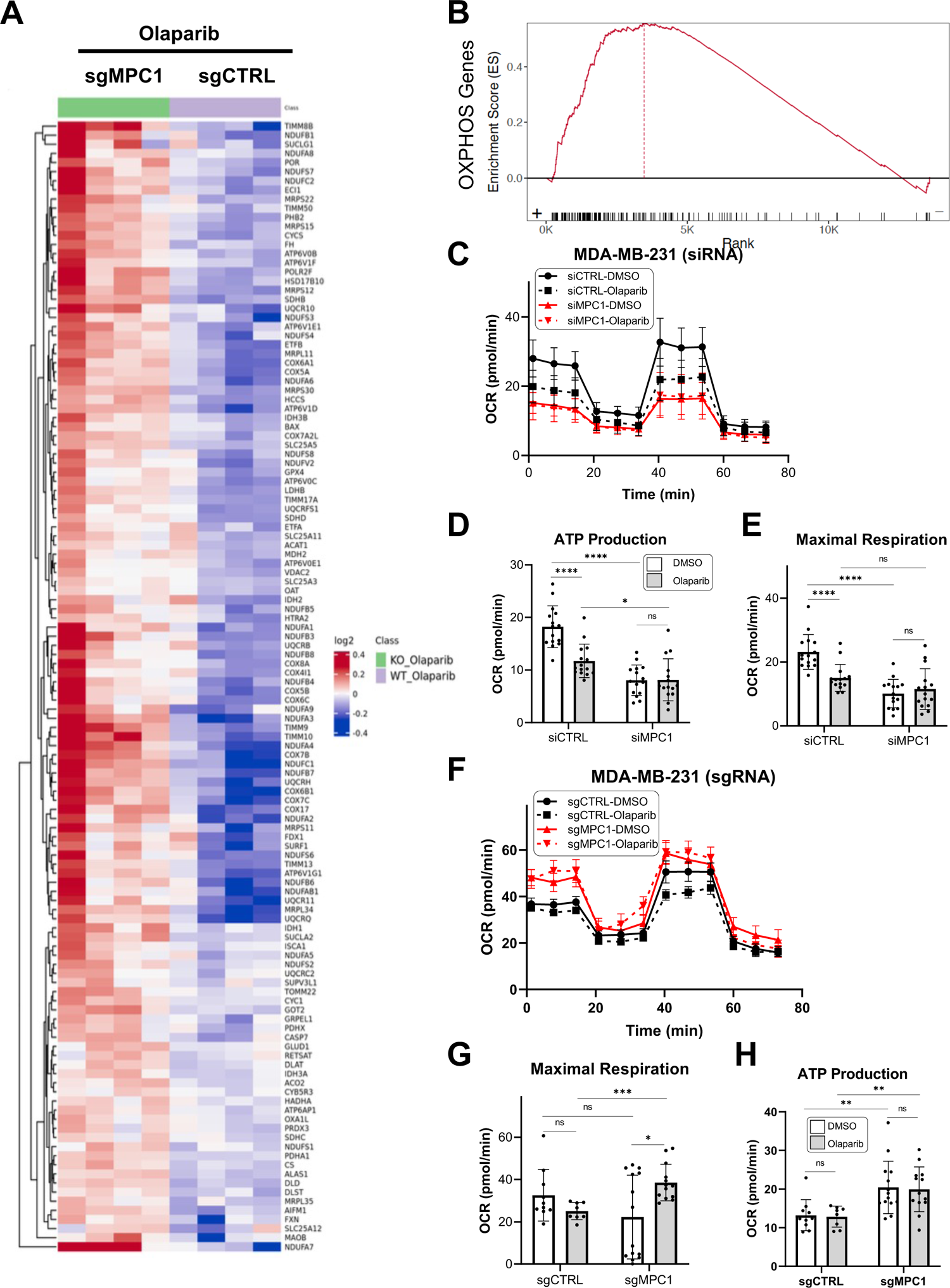
Oxidative phosphorylation is reactivated in MPC1-depleted MDA-MB-231 cells upon PARP inhibition with Olaparib. (A) Heatmap showing the 141 differentially expressed OXPHOS genes in control and MPC1-depleted cells (sgMPC1) following treatment with Olaparib (10μM) for 6 days. (B) Gene enrichment Score for 141 OXPHOS genes using *GSEA* analysis. Note that most OXPHOS genes (over 135 genes) are ranked (on the X axis) on the positive region of the enrichment (Y axis) while only a fraction of OXPHOS genes (less than 6 genes) ranked negatively. (C) *Seahorse* assay results showing oxygen consumption rate (OCR) comparing parental and MPC1-depleted MDA-MB-231 cells (siRNA targeting MPC1), treated with Olaparib (10μM) for 6 days. (D) ATP production of parental and MPC1-depleted MDA-MB-231 cells assayed in C. (E) Maximal respiration of parental and MPC1-depleted MDA-MB-231 cells assayed in C. (F) *Seahorse* assay results showing oxygen consumption rate (OCR) of MDA-MB-231 cells infected with sgCTRL or sgMPC1 (knockout pool) and treated with Olaparib (10μM) for 6 days. (G) ATP production of cells assayed by *Seahorse* in F. (H) Maximal respiration of cells assayed by *Seahorse* in F. Statistical significance was determined by two-tail unpaired student t-test. Data are represented as mean SD; n = 15. ns=not significant; *P < 0.05; ****P < 0.0001.

MPC1 transports pyruvate into the mitochondrial matrix, where pyruvate is oxidized to acetyl-CoA before its entry into the TCA cycle (Figure 5A). MPC1 expression is associated with the Warburg effect and cell survival^16–18^. To establish the extent to which MPC1 deletion and PARP inhibition affect energetic metabolism in breast cancer cells, we used a reversed-phase ion-pairing LC-MS^2^ assay to measure cell central carbon metabolites with a focus on the TCA metabolites. Our data revealed that PARP inhibition in 4T1 cells depleted of MPC1 led to a robust accumulation of Pyruvate (Figure 5B), Lactate and Acetyl CoA (Figure 5C, and D), and the TCA metabolites including Citrate, Cis-Aconitate, Succinate, Fumarate, and Malate (Figure 5E-J). The elevation in the TCA metabolites upon PARP inhibitor treatment ascertains the evidence of energic metabolism rewiring in breast cancer cells, findings which are consistent with the reactivation of Oxidative Phosphorylation (OXPHOS) and mitochondrial respiration in MPC1-depleted cells treated with Olaparib. These findings strongly suggest that the ability of breast cancer cells to resist PARP inhibitor treatment *in vivo* may reflect an intrinsic mechanism of metabolism rewiring, thus endowing tumor cells with a unique capacity to progress despite PARP inhibition. Other studies have demonstrated that oxidative phosphorylation is an essential process that drives cancer drug resistance and has a major influence on response to anticancer therapy ^19–22^. Taken together, these results suggest that a permanent loss of MPC1 sensitizes lung cancer cell lines to PARP inhibition but in contrast, endows breast cancer cell lines with the ability to rewire their oxidative phosphorylation capacity, thus, and overcome metabolic vulnerability (Figure 6). Whether this characteristic holds for most lung and breast cancer lines requires further study.

**Figure 5.**
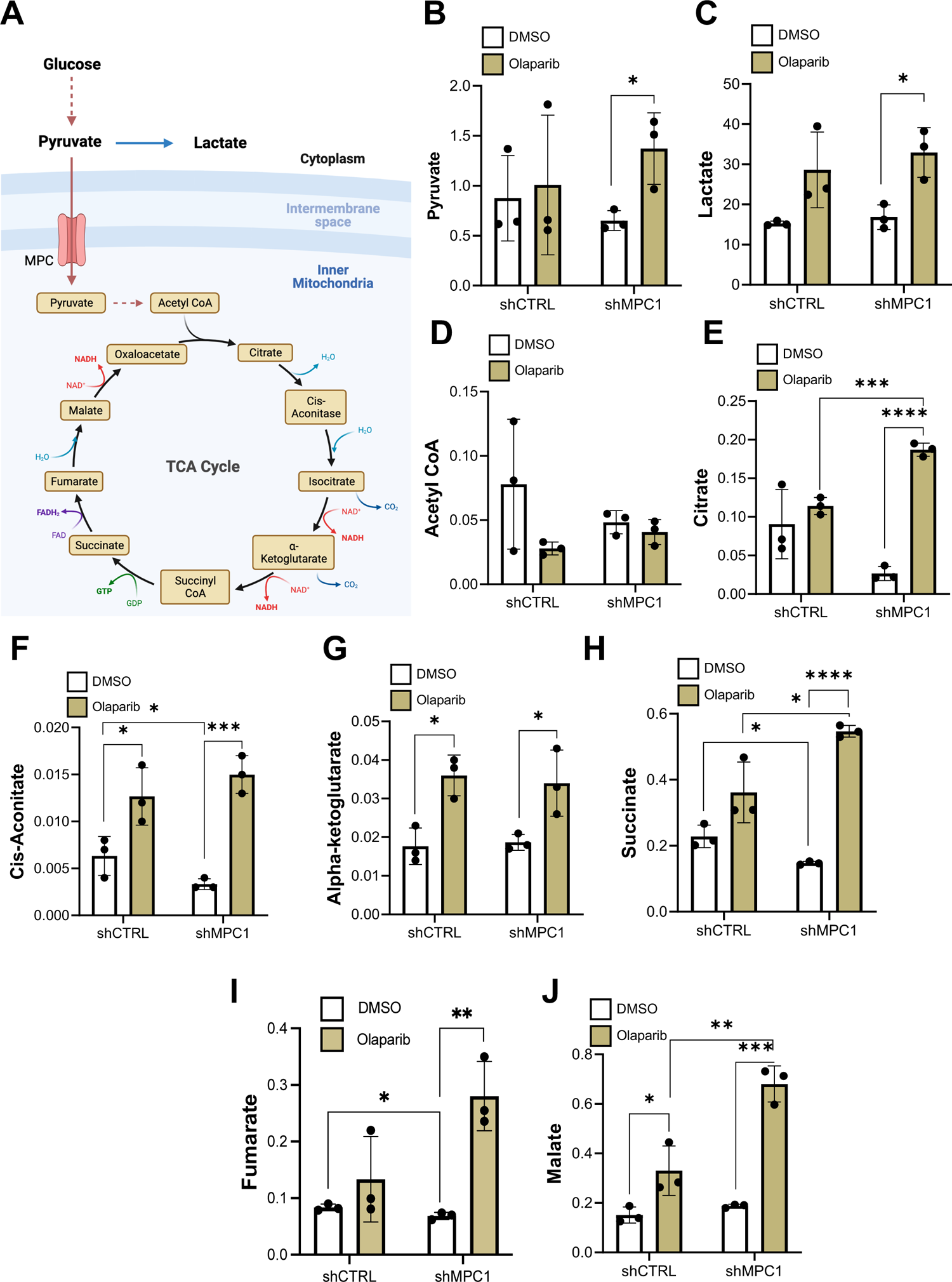
Effect of MPC1 knockdown on TCA cycle metabolites in 4T1 cells. (A) Schematics of the TCA cycle. (B) to (J) Selected TCA cycle metabolite levels (micromolar/million cells) of 4T1-shCTRL or 4T1-shMPC1 cells treated with DMSO or 10μM Olaparib for 6 days. Statistical significance was determined by two-tail unpaired student t-test. Data are represented as mean SD; n = 3. *P < 0.05; **P<0.01; ***P <0.001; ****P < 0.0001.

**Figure 6.**
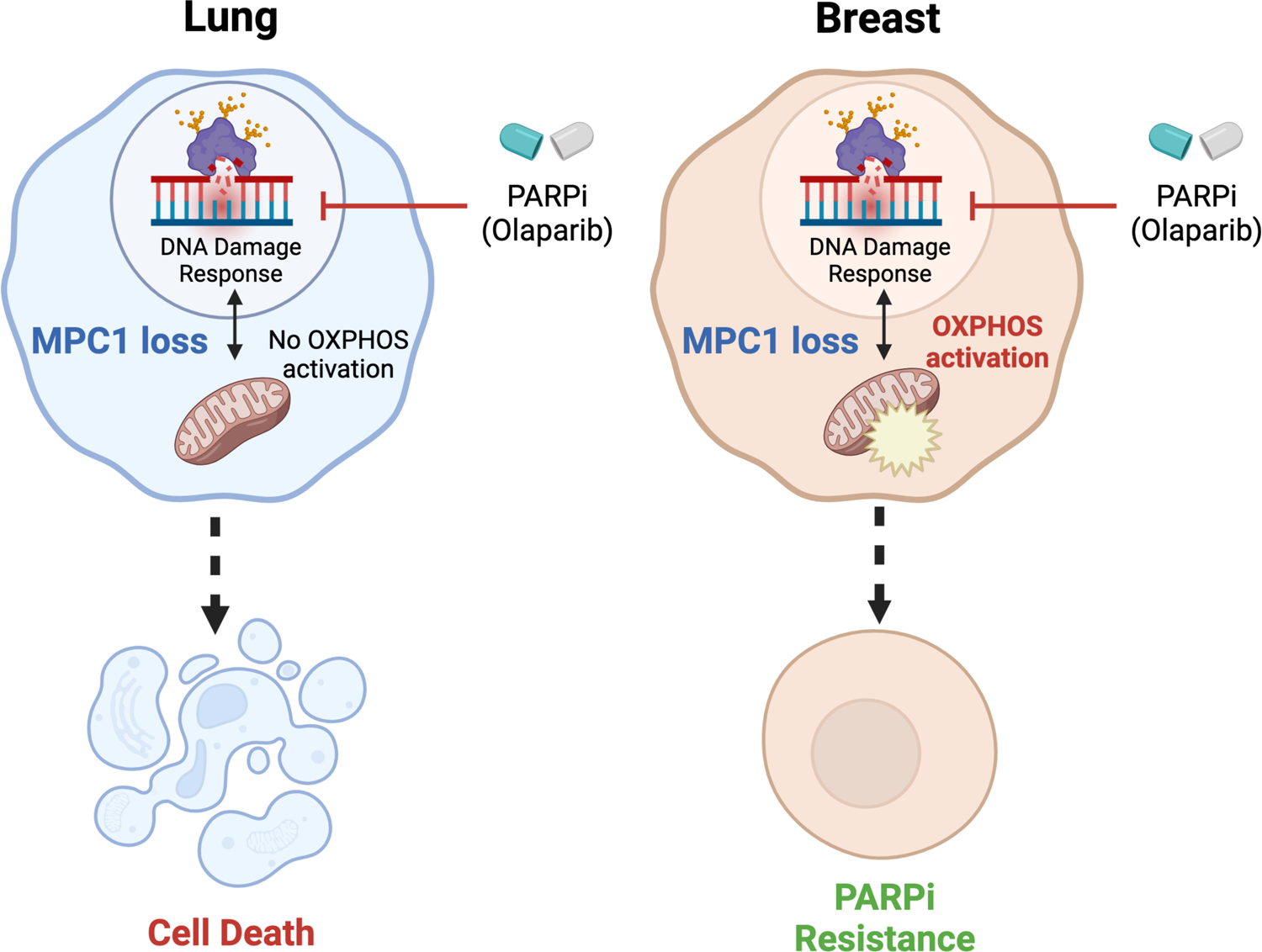
Model for the differential response to PARP inhibition in lung and breast cancer cells. Left, Lung cancer cell line. Right, Breast cancer cell line.

PARP inhibitors have entered broad clinical use, but their efficacy remains restricted to a subset of patients with HR gene mutations ^7, 23^. MPC1 is a robust metabolic sensor essential for pyruvate-driven mitochondrial respiration and cell survival ^16^. Our findings that MPC1 loss sensitizes lung cancer cells, but not breast cancer cells, to PARP inhibition *in vivo*, uncover a novel metabolic pathway that could be potentially exploited to improve PARPi efficacy. Understanding the mechanism underlying the regulation of mitochondrial homeostasis by MPC1 will provide a consolidated groundwork for elucidating the crosstalk between PARP-dependent DNA repair and mitochondrial functions, improving the clinical benefit of PARPi therapies. Finally, identifying MPC1 as a new metabolic player that influences PARPi treatment may unfold additional avenues for improving PARPi efficacy in cancer and benefitting a larger cohort of patients.

## Author contributions

T.F., R.C., Y.L., and U.W. designed the experiments. T.F., R.C., Y.L., H.L., T.A., X.X., W.R., O.W., M.B., T.J.M., and U.W. performed the analyses. T.F., R.C., and Y.L. performed the experiments. M.E.D, and P.Y. provided reagents and cell lines. T.F., R.C., Y.L., H.L., T.A., X.X., W.R., O.W., M.B., T.J.M., and M.E.D. edited the manuscript. U.W. wrote the manuscript with inputs from all authors.

## Acknowledgments

We thank Drs. William M. Bonner and Travis H. Stracker of the National Cancer Institute (NCI) for their thoughtful critiques and their help with reviewing this manuscript. We are indebted to Drs. Jing Huang and Hualong Yan for help with the *Seahorse* Analyzer. This work is supported by the Cancer Prevention and Research Institute of Texas grant (CPRIT RR190101), the Center for Cancer Research of the National Cancer Institute, and the Alfred P. Sloan Research Foundation (FG-2021-16433).

## Abbreviations

PARP: Poly (ADP-Ribose) Polymerase

PARPi: PARP Inhibitor

MPC1: Mitochondrial Pyruvate Carrier 1

PARylation: Poly ADP-ribosylation

HR: Homologous Recombination

NSCLC: Non-Small Cell Lung Cancer

OXPHOS: Oxidative Phosphorylation

OCR: Oxygen Consumption Rate

PTEN: Phosphatase and TENsin homolog

SiRNA: Small Interfering Ribonucleic Acid

SgRNA: Single guide Ribonucleic Acid

NAD: Nicotinamide Adenine Dinucleotide

TCA: Tricarboxylic acid

GSEA: Gene Set Enrichment Analysis

CRISPR: Clustered Regularly Interspaced Short Palindromic Repeats

KO: Knockout

NSG: NOD Scid Gamma

CCM: Central Carbon Metabolites

DMSO: Dimethyl Sulfoxide

GAPDH: Glyceraldehyde-3-phosphate Dehydrogenase

LC-MS: Liquid Chromatography – Mass Spectrometry

---

**Supplementary Figure 1.**
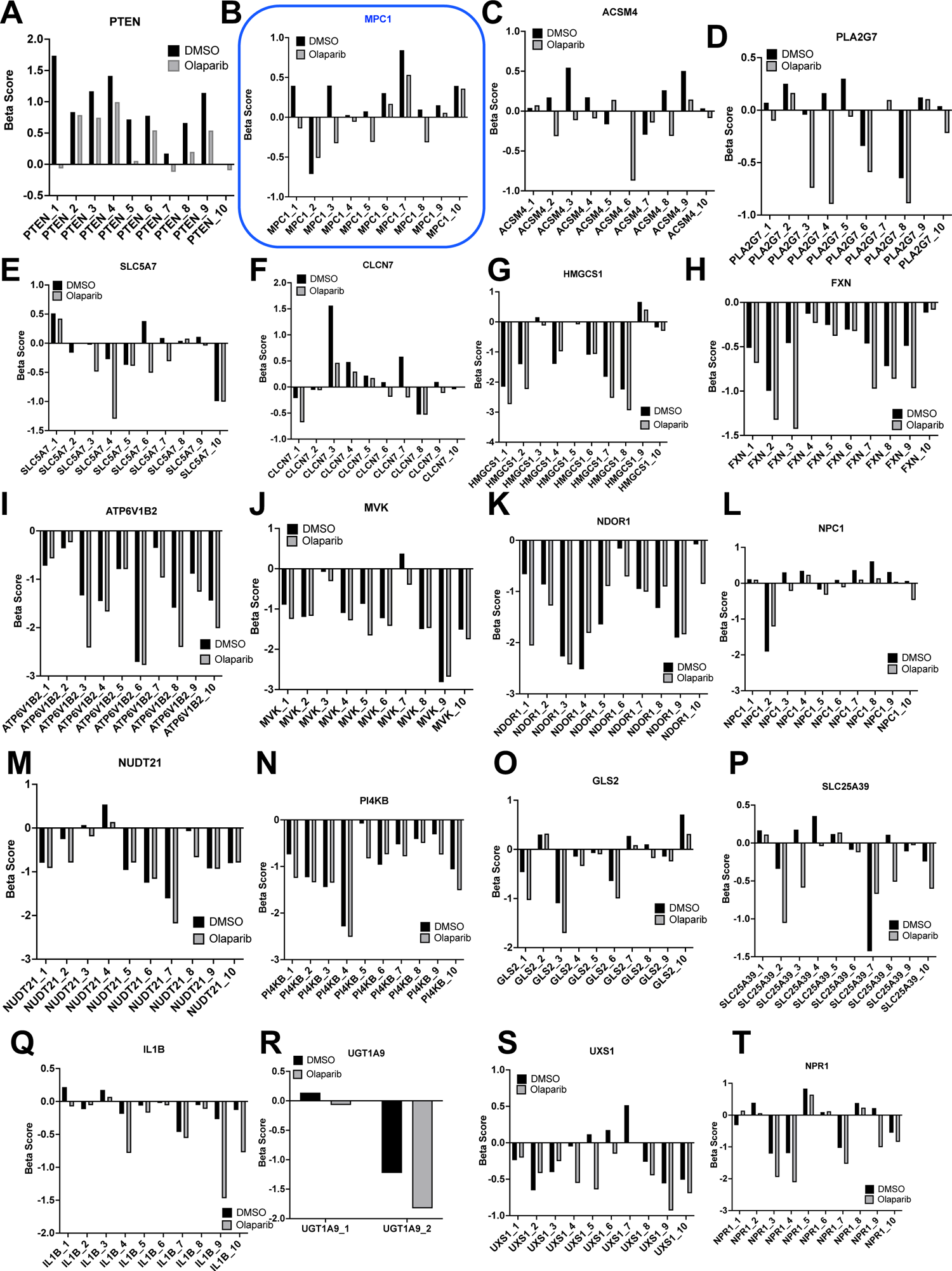
Distribution of sgRNAs targeting the top 20 genes identified by the CRISPR/Cas9 knockout screen. Changes in the abundance of the individual sgRNA guide in presence or absence of Olaparib.

**Supplementary Figure 2.**
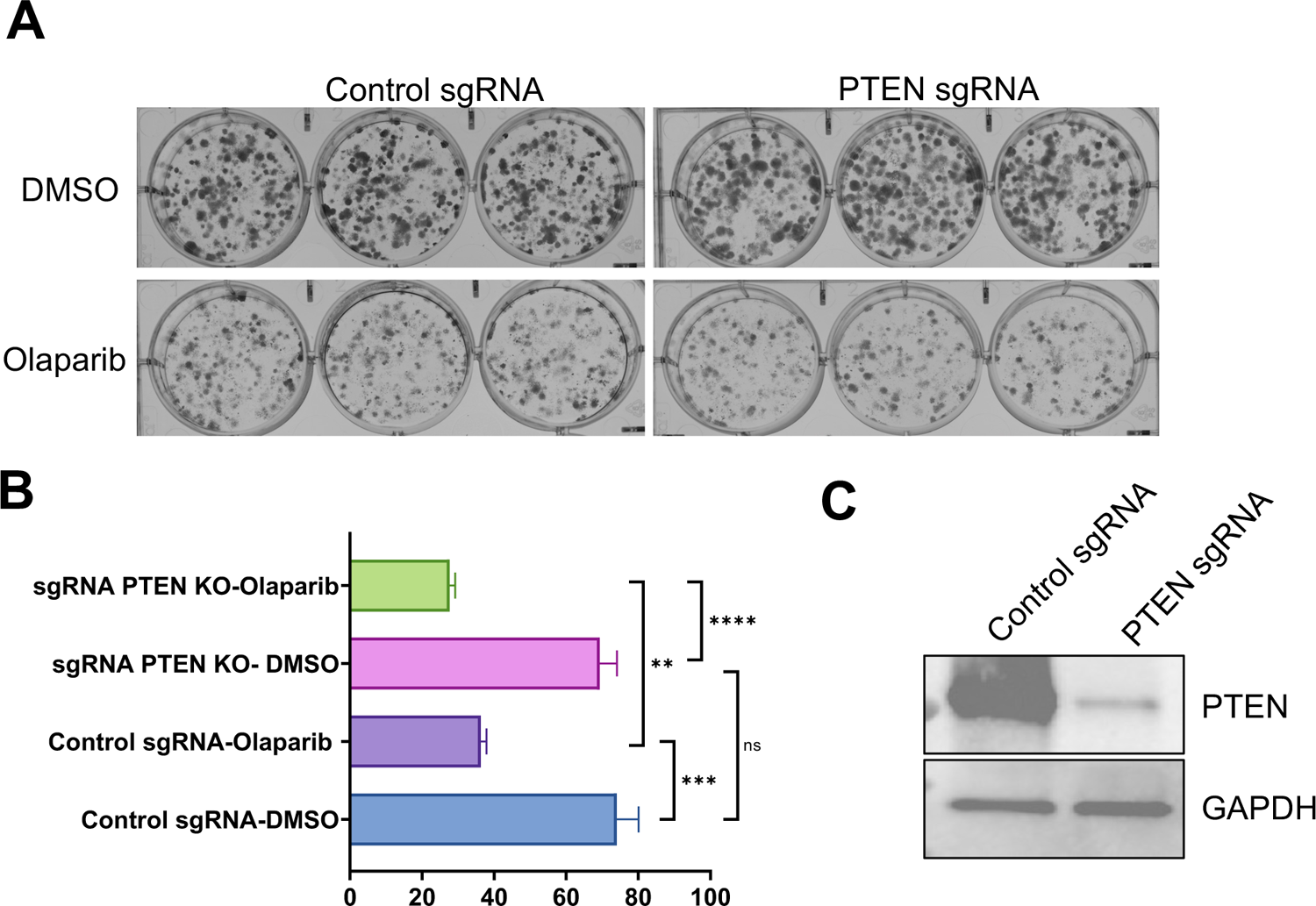
Loss of PTEN does not affect Olaparib sensitivity in MDA-MB-231 cells. (A) Clonogenic assays of control sgRNA or PTEN sgRNA MDA-MB-231 cells after treatment with DMSO or Olaparib (2.5 μM) for 10 days. (B) Quantification. (C) PTEN protein level is significantly reduced in sgPTEN infected MDA-MB-231 cells.

**Supplementary Figure 3.**
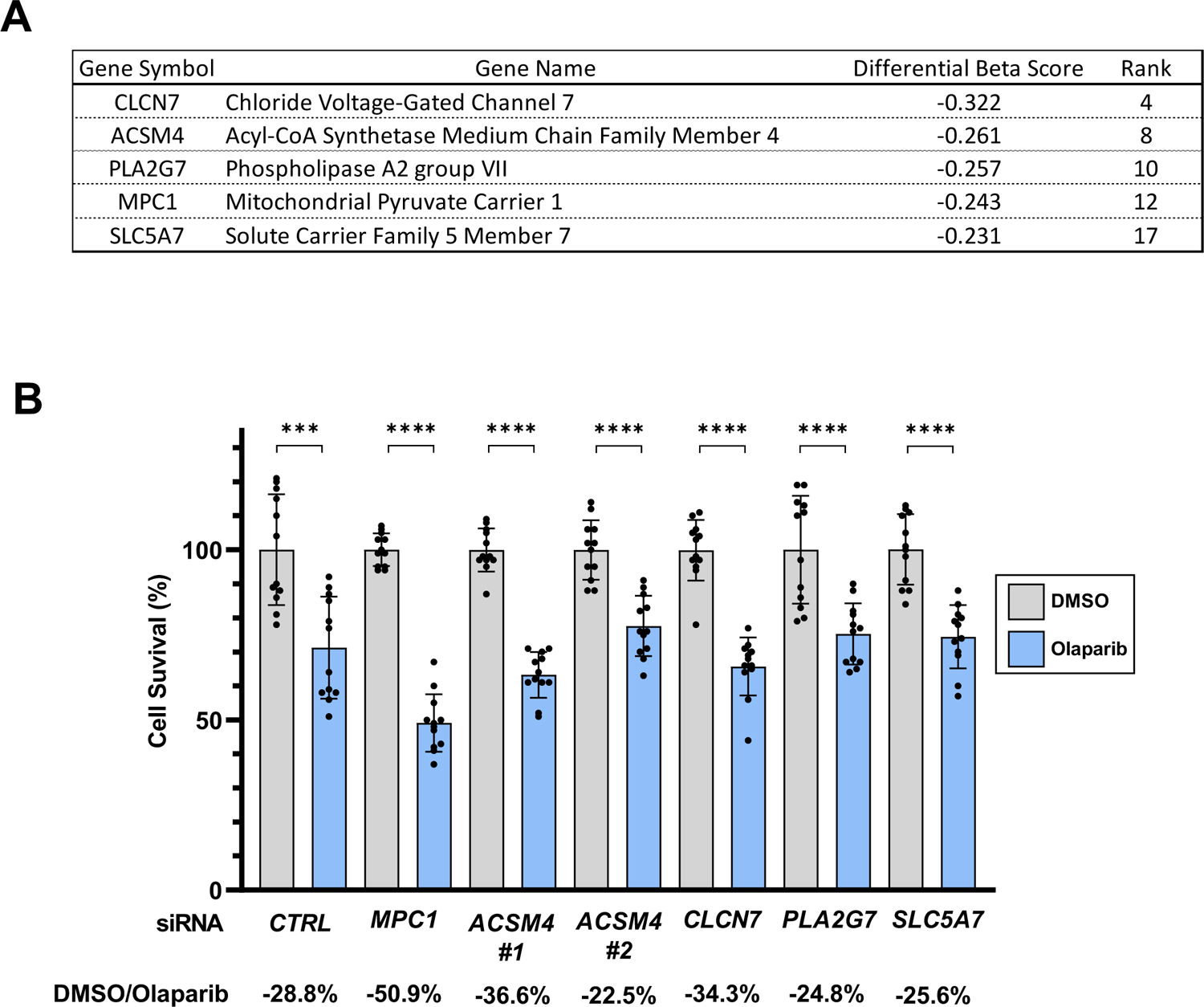
SiRNA screen to identify the most required gene for PARP inhibitor resistance. (A) List of genes with their beta-score and rank selected based among genes shown in Supplementary Figure 1 using sgRNAs distribution. (B) Survival of MDA-MB-231 cells transfected with control or siRNA targeting ACSM4, CLCN7, PLA2G7, MPC1, or SLC5A7, and treated with Olaparib (5 μM) for 6 days. Statistical significance was determined by two-tail unpaired student t-test. Data are represented as mean SD; n = 12. ***P <0.001; ****P < 0.0001.

**Supplementary Figure 4.**
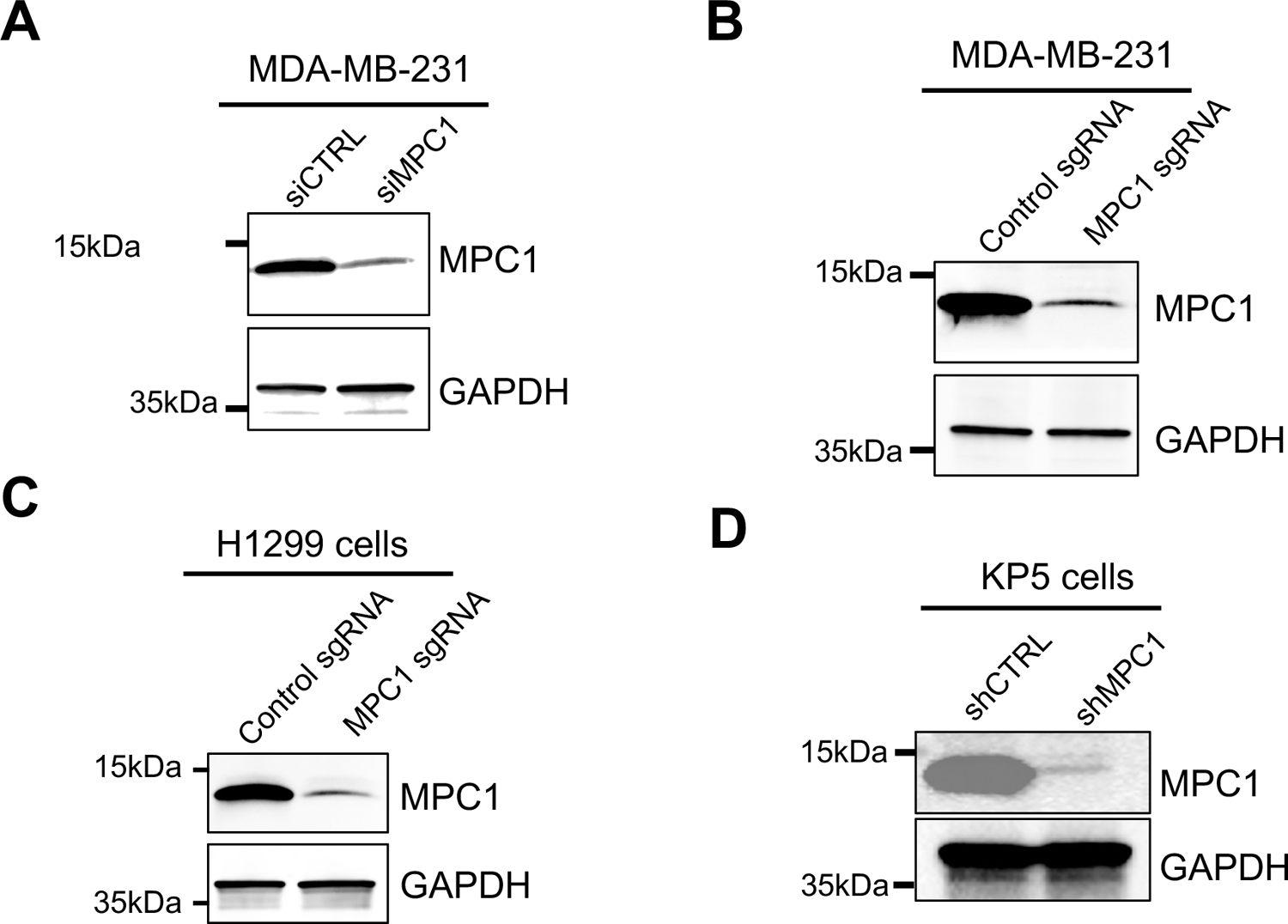
Validation of MPC1 downregulation by western blot. (A) Western blotting showing MDA-MB-231 cells transfected with siCTRL or siMPC1. Whole cell lysates were prepared 2 days post transfection. (B) Western blotting showing MDA-MB-231 cells infected with sgCTRL or sgMPC1. (C) Western blotting showing NCI H1299 cells infected with sCTRL or sgMPC1. (D) Western blotting of KP5 cells infected with shCTRL or shMPC1.

**Supplementary Figure 5.**
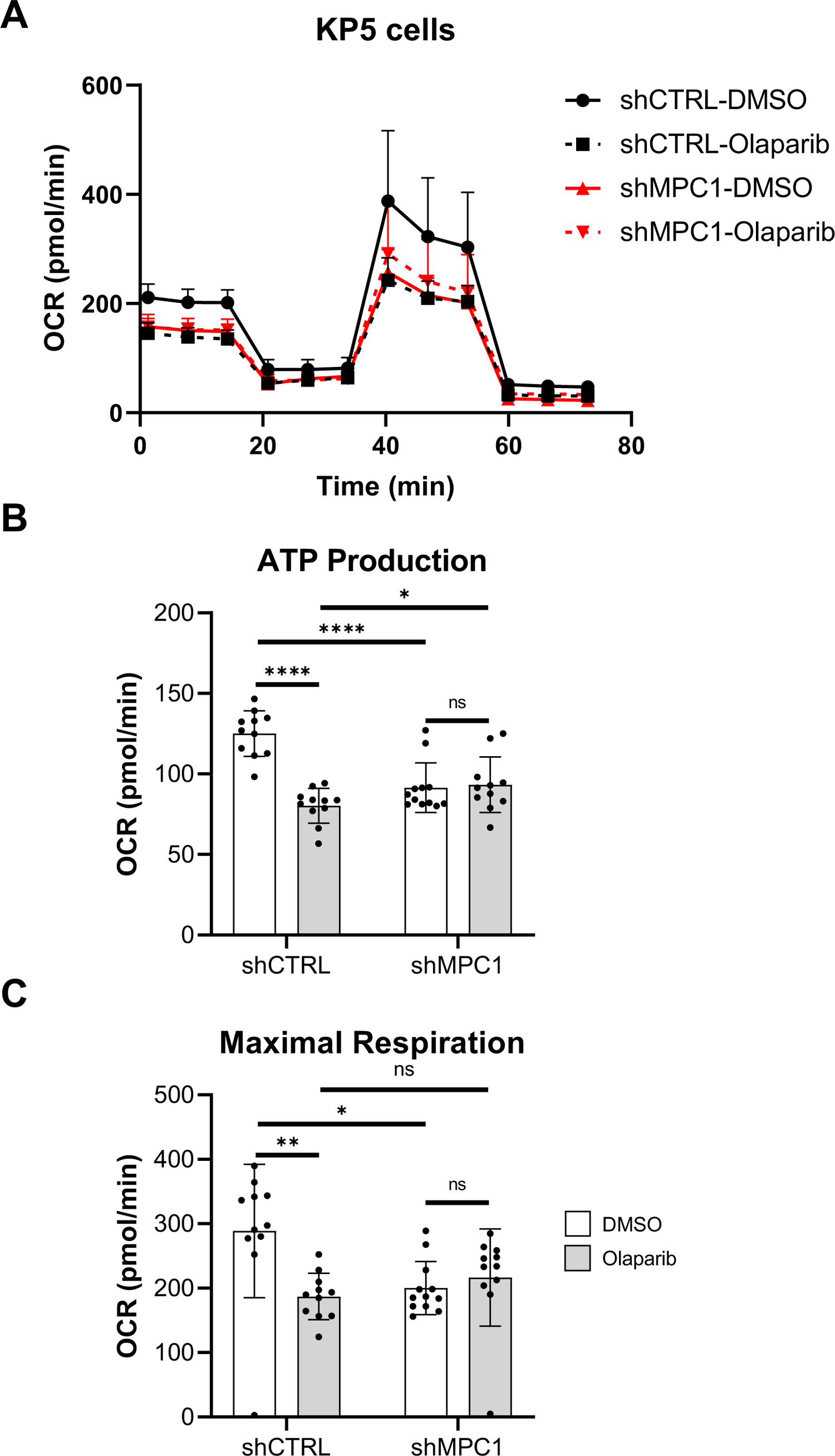
Seahorse assay comparing control and MPC1-depleted KP5 cells. (A) *Seahorse* assay to assess oxygen consumption rate (OCR) in KP5 shCTRL or KP5 shMPC1 cells treated with DMSO or Olaparib (10μM) for 6 days. (B) ATP production of cells assayed in A. (C) Maximal respiration of cells analyzed in A. Statistical significance was determined by two-tail unpaired student t-test. Data are represented as mean SD; n = 12. ns=not significant; *P < 0.05; **P< 0.001, ****P < 0.0001.

