## Supplementary material for "Metabolism-focused CRISPR screen unveils Mitochondrial Pyruvate Carrier 1 as a critical driver for PARP inhibitor resistance in lung cancer": Table 1

**Table1. Summary of CCM Targets and method**

| **Abbreviation** | **CCM** | **SI-CCM** | **ESI** | **Derivative** | **Native** |
| --- | --- | --- | --- | --- | --- |
| *TCA Cycle* |  |  |  |  |  |
| PYR | pyruvate; | 13C3_LAC | − | No | TBAA |
| LAC | Lactate | 13C3_LAC | − | No | TBAA |
| AcCoA | acetyl-coenzyme A; | 13C4_SUC | − | No | TBAA |
| CIT | citrate; | 13C4_SUC | − | No | TBAA |
| ACT | cis-aconitate; | 13C4_SUC | − | No | TBAA |
| AKG | 2-oxoglutarate; | 13C4_SUC | − | No | TBAA |
| SUC | succinate; | 13C4_SUC | − | No | TBAA |
| FUM | fumarate; | 13C4_SUC | − | No | TBAA |
| MAL | malate; | 13C4_SUC | − | No | TBAA |
| 2HG | 2-hydroxyglutaric acid | 13C4_SUC | − | No | TBAA |
| *Glycolysis* |  |  |  |  |  |
| G6P | glucose-6-phosphate; | 13C6_G6P | − | No | TBAA |
| F6P | fructose-6-phosphate; | 13C6_G6P | − | No | TBAA |
| FBP | fructose-1,6-diphosphate; | 13C6_FBP | − | No | TBAA |
| DHAP | dihydroxy-acetone-phosphate; | 13C6_G6P | − | No | TBAA |
| GAP | glyceraldehyde-3-phosphate; | 13C6_G6P | − | No | TBAA |
| 3PG | 3-phosphoglycerate; | 13C6_FBP | − | No | TBAA |
| PEP | phosphoenolpyruvate; | 13C6_FBP | − | No | TBAA |
| *Pentose Phosphate Pathway* |  |  |  |  |  |
| 6PG | 6-phosphogluconate; | 13C6_FBP | − | No | TBAA |
| R5P | ribose-5-phosphate; | 13C6_FBP | − | No | TBAA |
| Ru5P | ribulose 5-phosphate | 13C6_FBP | − | No | TBAA |
| XYLU5P | xylulose-5-phosphate; | 13C6_FBP | − | No | TBAA |
| S7P | sedoheptulose-7-phosphate; | 13C6_FBP | − | No | TBAA |
| *OxiPhos and Redox* |  |  |  |  |  |
| AMP | adenosine monophosphate | 13C4_SUC | − | No | TBAA |
| ADP | adenosine diphosphate | 13C4_SUC | − | No | TBAA |
| ATP | adenosine triphosphate | 13C4_SUC | − | No | TBAA |
| GTP | guanosine triphosphate | 13C4_SUC | − | No | TBAA |
| cAMP | adenosine 3′,5′-cyclic monophosphate | 13C4_SUC | − | No | TBAA |
| cGMP | guanosine 3′,5′-cyclic monophosphate | 13C4_SUC | − | No | TBAA |
| NAD | nicotinamide adenine dinucleotide | 13C4_SUC | − | No | TBAA |
| NADH | nicotinamide adenine dinucleotide (reduced) | 13C4_SUC | − | No | TBAA |
| NADP | nicotinamide adenine dinucleotide phosphate | 13C4_SUC | − | No | TBAA |
| NADPH | nicotinamide adenine dinucleotide phosphate (reduced) | 13C4_SUC | − | No | TBAA |
| FAD | flavin adenine dinucleotide | 13C4_SUC | − | No | TBAA |
| GSH | glutathione | 13C4_SUC | − | No | TBAA |
